# Regulation of the Promoter for Capsular Polysaccharide Synthesis in *Neisseria meningitidis* Serogroup B by HTH_XRE Family Transcription Factor

**DOI:** 10.1101/2024.10.23.619888

**Authors:** Iyinoluwa Sofowora, Pumtiwitt McCarthy, James Wachira

## Abstract

The capsular polysaccharide synthesis (*cps*) locus of *Neisseria meningitidis* is implicated in invasive meningococcal disease. The synthesis (*synABCD*) and transport (*ctrABCD*) operons are transcribed in opposite directions from a common intergenic region and expression is negatively regulated by the bacterial two-component system *misR/misS* and thermosensitive RNA folding. However, these mechanisms do not fully explain the stationary phase responses and the cis-acting elements remain to be fully characterized. Using GFP reporter gene and site-directed mutagenesis, cis-regulatory elements in the 134-bp intergenic region, NmIR, were investigated. While confirming a known RpoD promoter, an additional potential promoter element and putative binding sites for the transcription factors *fis and lexA* were identified through sequence analysis. Deletion of the putative LexA binding site led to an increase in GFP fluorescence. The *N. meningitidis genome carries only one lexA homolog, the* Helix-Turn-Helix regulator XRE family member (GenBank-NMB0910, HTH_XRE). Trans-complementation of the NmIR-GFP reporter with the *N. meningitidis* HTH_XRE expression plasmid led to increased fluorescence. Trans-complementation with either *misR/misS* or *nusG* decreased reporter gene expression. Consistent with previous reports, deletion of the RpoD promoter reduced expression by 50%, suggesting a redundancy of promoter elements in the intergenic region. Thus, the results confirm the functioning of an exogenous *N. meningitidis* CPS synthesis promoter in *E. coli* and demonstrate its regulation through trans-complementation by *misR/misS,* HTH_XRE, and *nusG*.

**Importance:** Pathogenic *Neisseria meningitidis*, a causal agent of bacterial meningitis, secretes capsular polysaccharides of different compositions that differentiate the serogroups. Since the capsule is an important virulence factor that determines adhesion to epithelia and ability to invade tissues, there is need to understand the underlying mechanisms for its expression. Furthermore, bacterial polysaccharides are potential sources of novel biomaterials. The expression of the capsule production genes is regulated, and this study reveals a mechanism involving a transcription factor, HTH_XRE, whose function in *Neisseria meningitidis* is not known. It extends the understanding of capsular expression regulation by identifying other control elements in the promoter region. The results will have applications in optimizing biomaterial production or in developing therapeutic interventions.

## Introduction

*Neisseria meningitidis* is a gram-negative bacterial pathogen responsible for endemic meningitis. The capsule plays a significant role in the pathogenesis of the disease and has been the target for meningococcal vaccines (1, 2). Among the twelve serogroups of *N. meningitidis* defined by capsular polysaccharides, six serotypes (A, B, C, W, X, and Y) are known to cause invasive diseases (2, 3). The genomic organization of the *cps* loci in *N. meningitidis* has been well studied and comprises three regions: regions A, B, and C (4). Region A encodes three s*ynABC genes,* which are required for sialic acid synthesis, and the serogroup-specific gene *siaD,* encoding polysialyltransferase. Region C, which codes for the capsule transport operon, is highly conserved among all serogroups and comprises four genes, ctrA-D, which are responsible for the transport of capsule polymers to the bacterial surface. Region B contains genes for polysaccharide translocation (5, 6). Except in serogroups I and K, where the operons are transcribed in the same direction, the promoters for both the synthesis and transport operons are contained in a 134-bp intergenic region (NmIR) (5). Capsular expression is regulated by several mechanisms, including transcriptional and translational mechanisms (5). However, the control elements have not been fully identified.

At the transcription level, von Loewenich and coworkers identified an UP-like element and an untranslated region as key determinants of capsule synthesis (7). The deletion of the UP-like element reduced reporter gene expression (7). The sequential deletion of the -13 to +53 region led to a total loss of expression (7). Deletion of the +13 to +103 region led to an increase in expression of 11-fold, suggesting the presence of an inhibitory sequence (7). However, the mechanism of the function and physiological role of this region remain unknown, as environmental conditions such as temperature, glucose concentration, and iron concentration do not affect capsule production (7). However, the folding of the transcript to include the direct repeat and a segment that includes the ribosome binding site (RBS) is proposed to form a stable hairpin that regulates translation in some strains in a temperature sensitive manner (5).

The UP-like element is juxtaposed to the -35 promoter box on the 5’ end, and another study identified a contact-regulated gene A (*crgA*)/*LysR* transcription factor-binding site on the 3’ side that overlaps this element (8). This CrgA binding site is a negative regulatory element that functions to downregulate *syn* genes during adhesion to epithelial cells, but not during growth in suspension (8). The *crg*A gene is upregulated during coculture with epithelial cells together with the regulatory proteins *misR*/*phoP*, a two-component global regulatory system that is also implicated in the regulation of capsule expression (9). Coculture with epithelial cells, which promotes adhesion, is accompanied by downregulation of genes involved capsule production and other virulence factors in association with the upregulation of MisR (9). MisR/MisS was shown to regulate capsule expression and to bind directly to NmIR (10). Mutation of the *misR/misS* two-component regulatory system is associated with increased capsule expression, and MisR binds to DNA encompassing NmIR and the 5’-coding region of *cssA* (5, 10).

Nevertheless, the MisR binding element and signaling mechanisms upstream of MisS are not known. MisR is the DNA binding partner of the MisR/MisS two-component system, and the consensus binding site was identified as KWWWTGTAARGNNWH (11). MisR was demonstrated to bind to fourteen promoters within the genome, including the promoters of three transcription factors that carry the helix-turn-helix (HTH) DNA binding domain (5, 11, 12).

The expression of pathogenesis factors, including the capsule, was shown to be regulated by temperature through thermosensitive 5’-UTRs (13). Folding of the RNA transcripts suggested that the thermosensor functions by sequestering the ribosome binding site within secondary structure elements, and mutants with deletions in the direct repeats expressed elevated levels of CssA protein, whereas the levels of *cssA* mRNA were unaffected (5). In this study, a negative regulatory element with sequence homology to LexA binding sites was identified in the direct repeat sequence. HTH_XRE, an *N. meningitidis* LexA homolog, upregulated NmIR-dependent transcription.

## RESULTS

### Identification of putative binding sites and promoters

Softberry BPROM software (14) analysis of NmIR identified putative binding sites for the TFs *fis and lexA* and three *rpoD* sites *(rpoD 16, rpoD 17, and rpoD 18)* (Table 1 and Fig. 1). Two promoters were identified: one adjacent to the up-like element and one at the direct repeat region. The predicted promoter consensus -10 box (TTATATACT) and a -35 box (TTTCCA) were identified upstream of the *synA* gene (GenBank accession number: X78068.1). The direct repeat within the NmIR was predicted to be important for transcription via BPROM software.

**Table 1.** BPROM Promoter Prediction. A 249-sequence encompassing the initial coding regions of *ctrA* and *synA* was used to detect promoter elements.

| TF | Score | Sequence | Location Relative to the Initiation<br>Codon |
| --- | --- | --- | --- |
| *rpoD16* | 9 | GAGTATAA | -59 |
| *rpoD18* | 10 | CTTATACT | -28 |
| *fis* | 18 | TATACTTA | -26 |
| *lexA* | 14 | TATATATA | -20 |
| *lexA* | 14 | TATATATA | -18 |
| *lexA* | 14 | TATATATA | -16 |
| *lexA* | 12 | ATATATAC | -15 |
| *lexA* | 18 | TATATACT | -14 |
| *fis* | 18 | TATACTTA | 12 |
| *rpoD17* | 11 | TATAGATT | -6 |

**Figure 1.**
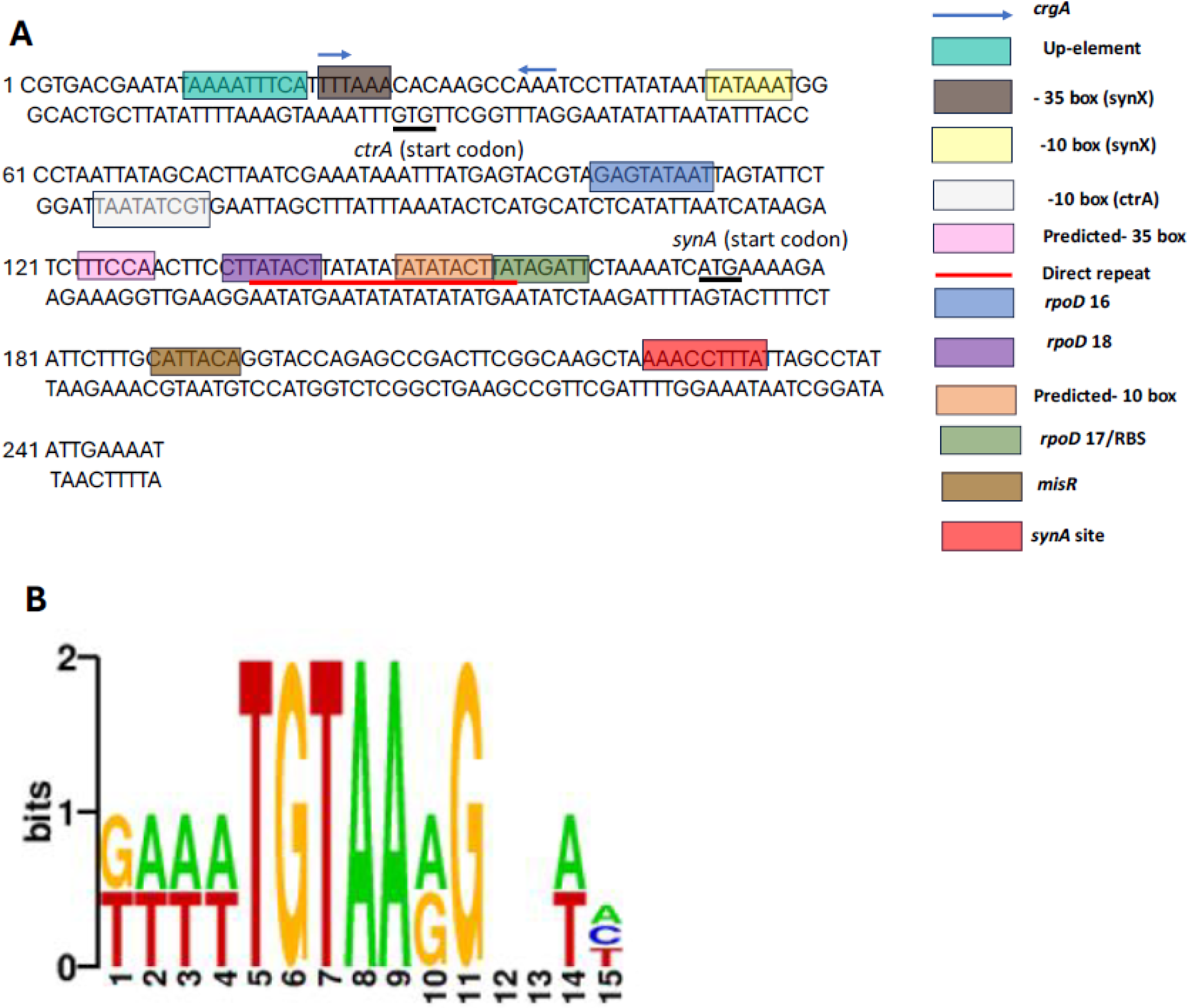
Regulatory elements within NmIR. (A) Key regulatory elements within the intergenic region upstream of the synA start codon are highlighted in colored boxes. The control elements include the promoter sequences (−10 box and -35 box), the up-like element, and the TF binding sites. (B) Consensus sequence of the misR binding site in NmIR.

*MisR/misS* has been reported to be a negative regulator of capsule expression in *N. meningitidis* (15, 16). Using FIMO software, we identified a putative binding site for *misR* within the *synA* gene, characterized by a consensus sequence of GAAATGTAAGGGGAA with a P value of 9.99e-02 (Fig. 1). Previous reports have demonstrated the binding of *misR* to a 600 bp DNA fragment encompassing the intergenic region and including the flanking coding regions of *synA* (10).

### Functional analysis of NmIR in *E. coli*

The studies were conducted in *E. coli. N. meningitidis* capsular genes can be exogenously expressed in *E. coli* from their natural promoter, with greater expression at the stationary phase (17). This was recapitulated by the NmIR-GFP-pRSET construct (Fig.2). NmIR-GFP-pRSET is fluorescent in *E. coli* DH5α and BL21(DE3) strains, whereas the pRSET vector alone without NmIR is only fluorescent in BL21(DE3) strains, whereby it is expressed from the T7 promoter. Elevated expression is observed as cultures transition to the stationary phase (Fig. 2).

**Figure 2.**
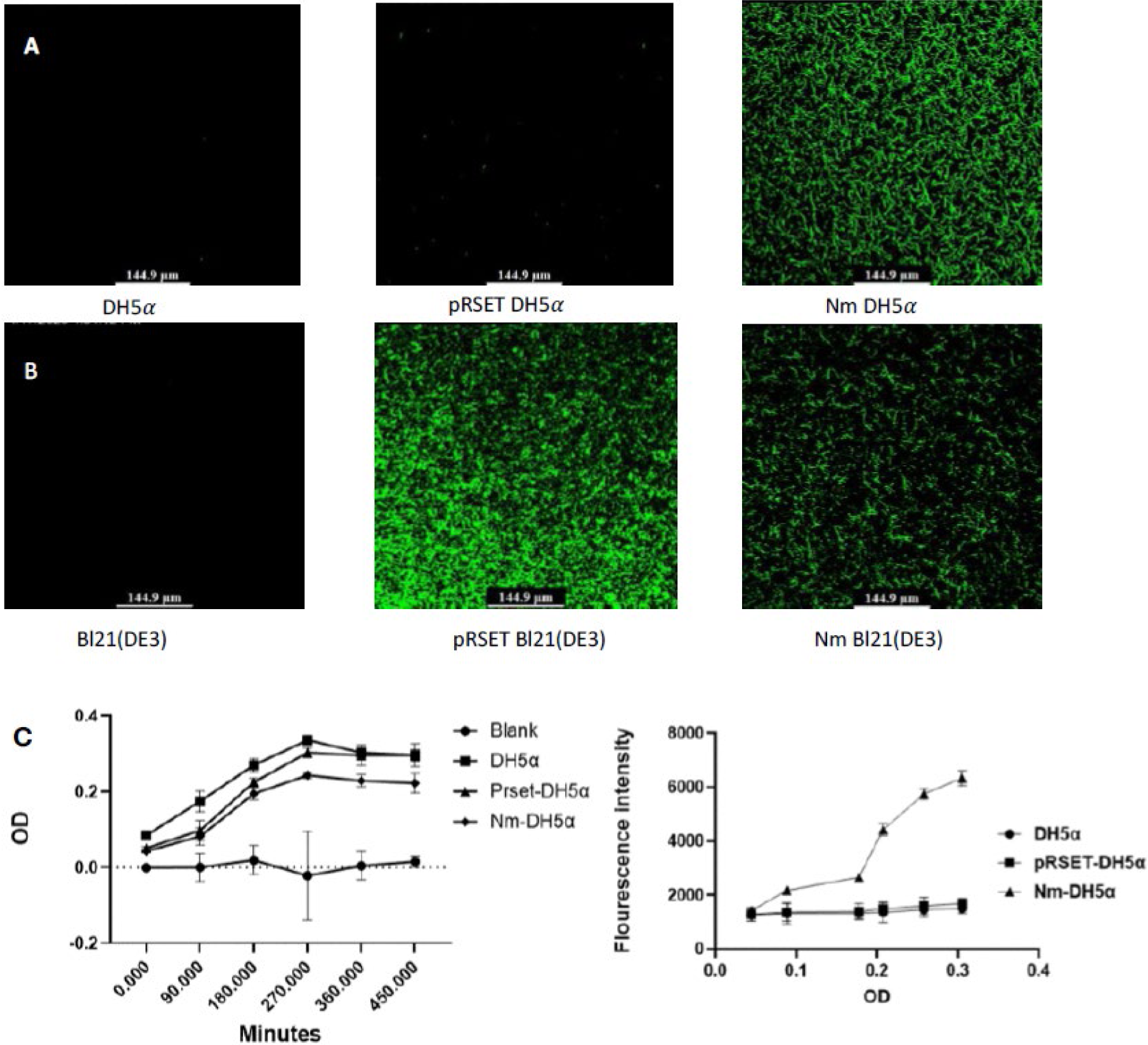
Functional analysis of GFP expression from NmIR. (A) DH5α, pRSET-EmGFP and NmIR-GFP-pRSET constructs expressed in DH5α. (B) BL21(DE3), pRSET-EmGFP and NmIR-GFP-pRSET constructs were transformed into BL21(DE3) cells. No fluorescence was detected in the pRSET-EmGFP expressed in DH5α, while fluorescence was detected in BL21(DE3) cells. In contrast, the NmIR-GFP-pRSET construct is fluorescent when expressed in *E. coli* DH5α and BL21(DE3) strains. (C) Expression curve of Nm as a function of the growth stage. The results revealed that there was more expression at the stationary phase.

### Role of predicted control elements in capsular expression

Deletion and GFP fluorescence quantification of the -10 promoter, up-like element, up-like element&-35 box, *rpoD* 16, and *rpoD* 17 regions of NmIR resulted in an approximately 50% reduction in GFP expression. Deletion of the *lexA* region is also predicted to encode a -10 box, leading to a 47% increase in capsular production. Deletion of the first repeat sequence led to a 15% decrease in GFP expression. Deletion of the *rpoD* 18 element and the direct repeat had no significant effect on capsular expression, as shown in Table 2 and Figure 3.

**Table 2.** Cis-acting elements regulating the *synABCD* operon. The predicted elements were deleted though site-directed mutagenesis and GFP fluorescence intensity determined. Image analysis was conducted with ImageJ (*43*).

| **NmIR**<br>**mutation** | **Mean Fluorescence**<br>**Intensity (arbitrary**<br>**units)** | **% WT** | **Standard**<br>**deviation** | **P-value** |
| --- | --- | --- | --- | --- |
| WT | 2.61 | 100 | 0.43 |  |
| DR | 2.61 | 100.02 | 0.36 | >0.9999 |
| 1st DR | 1.96 | 75.122 | 0.25 | <0.0001 |
| 10 Box | 1.13 | 43.54 | 0.18 | <0.0001 |
| *lexA* | 3.84 | 147.32 | 0.61 | <0.0001 |
| *rpoD* 16 | 1.48 | 56.84 | 0.30 | <0.0001 |
| *rpoD* 17/RBS | 1.13 | 43.53 | 0.11 | <0.0001 |
| *rpoD* 18 | 2.52 | 96.53 | 0.24 | 0.9785 |
| Up | 1.17 | 44.90 | 0.13 | <0.0001 |
| Up& -35 | 1.27 | 48.90 | 0.27 | <0.0001 |

**Figure 3.**
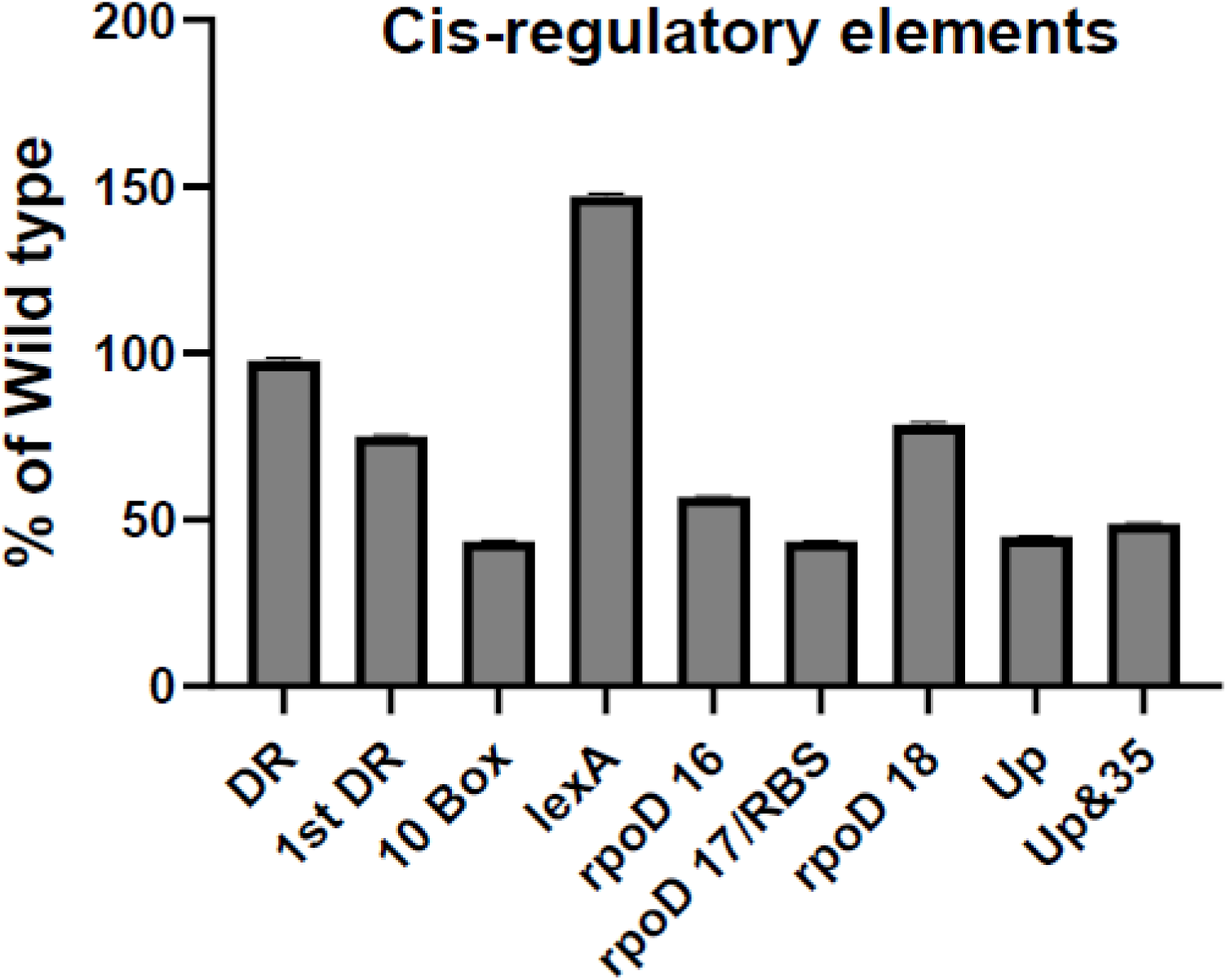
Percentage of wild-type expression of regulatory elements. Deletion of cis-regulatory elements (−10 promoter regions, rpoD 16, rpoD 17 binding sites, and Up-like elements) led to a 50% reduction in capsular expression, whereas lexA (the predicted -10 promoter) led to a 47% increase in capsular expression.

### Role of RNA secondary structure

Among the three mechanisms of capsule expression that impinge on the NmIR are translational control-mediated secondary structure formation in the 5’UTR (5). The deletion mutations were analyzed with RNAfold (18). The *lexA* deletion led to apparent destabilization of the secondary structure based on the free energy (−15.72 kcal/mol for the wild type as opposed to -10.07 kcal/mol for the mutant) (Fig. 4).

**Figure 4.**
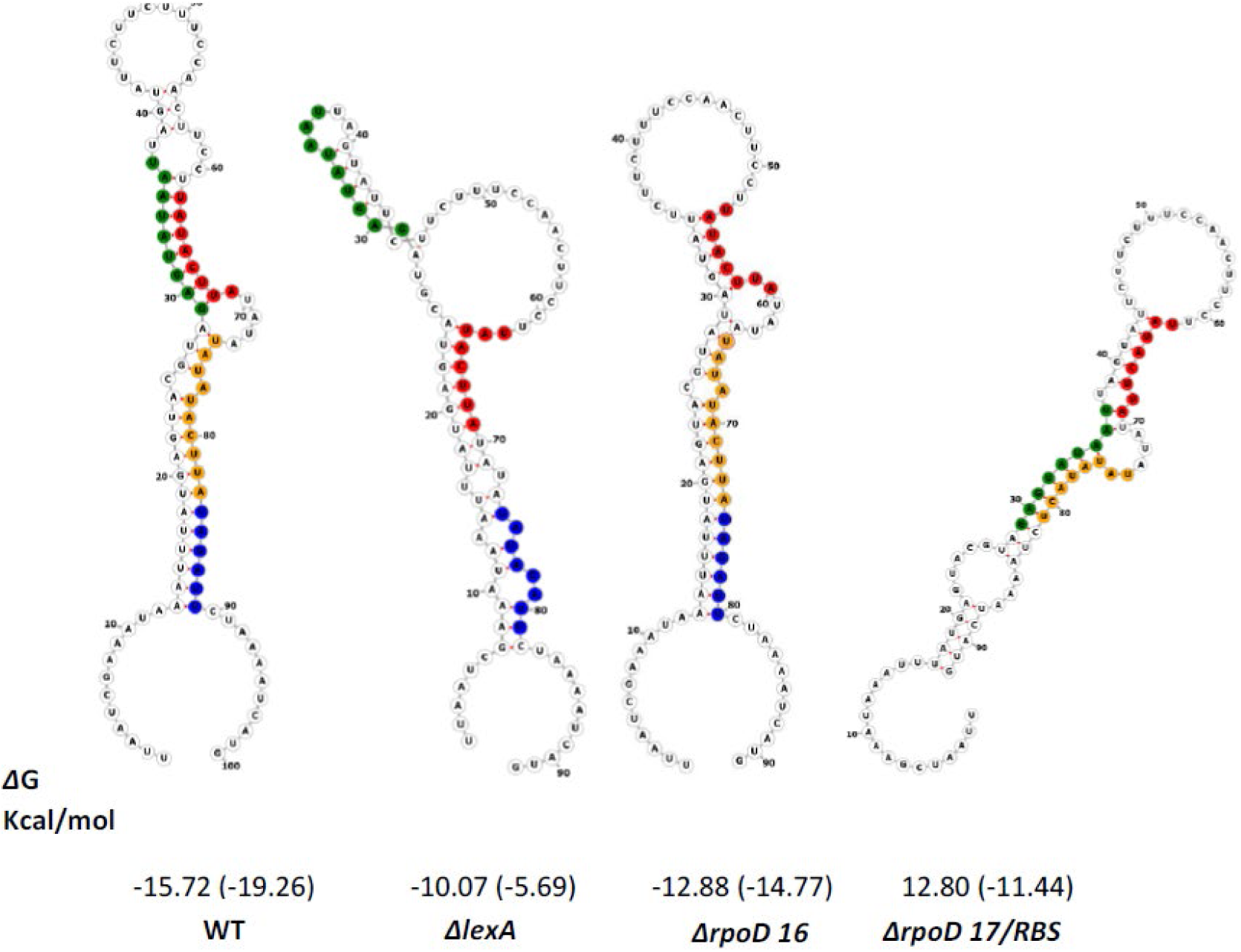
Predicted secondary structures of the RNA sequences from NmIR, wild type (WT), ΔlexA mutant, ΔrpoD 16 mutant, and ΔrpoD 17/RBS mutant. The colored regions highlight specific sequence elements, including the first copy of the direct repeat (red), the second copy of the direct repeat/lexA site (orange), the rpoD 16 site (green), and the ribosomal binding site, which overlaps with the rpoD 17 site (blue). The secondary structures were predicted via RNAfold, with free energy changes (ΔG) in kcal/mol shown below each structure.

### Cis-regulatory elements for capsule expression in *N. meningitidis B* and related bacteria that use ABC transport-dependent pathways

The intergenic region in sialic acid-producing *N. meningitidis* serogroups (B, C, W, and Y) is conserved, with differences observed only in the repeat sequence (TATACTATACTTA), which varies between serogroups. Analysis of similar regions in *E. coli* and *Haemophilus influenzae* (Fig. 5) revealed the conservation of AATAAAAG and TATATACT. These sequences are putative binding sites for *phoB, rpoD,* and *lexA*. This finding is consistent with the data showing the loss of expression of GFP at the mutated *rpoD* site and increased expression of GFP at the mutated *lexA* site.

**Figure 5.**
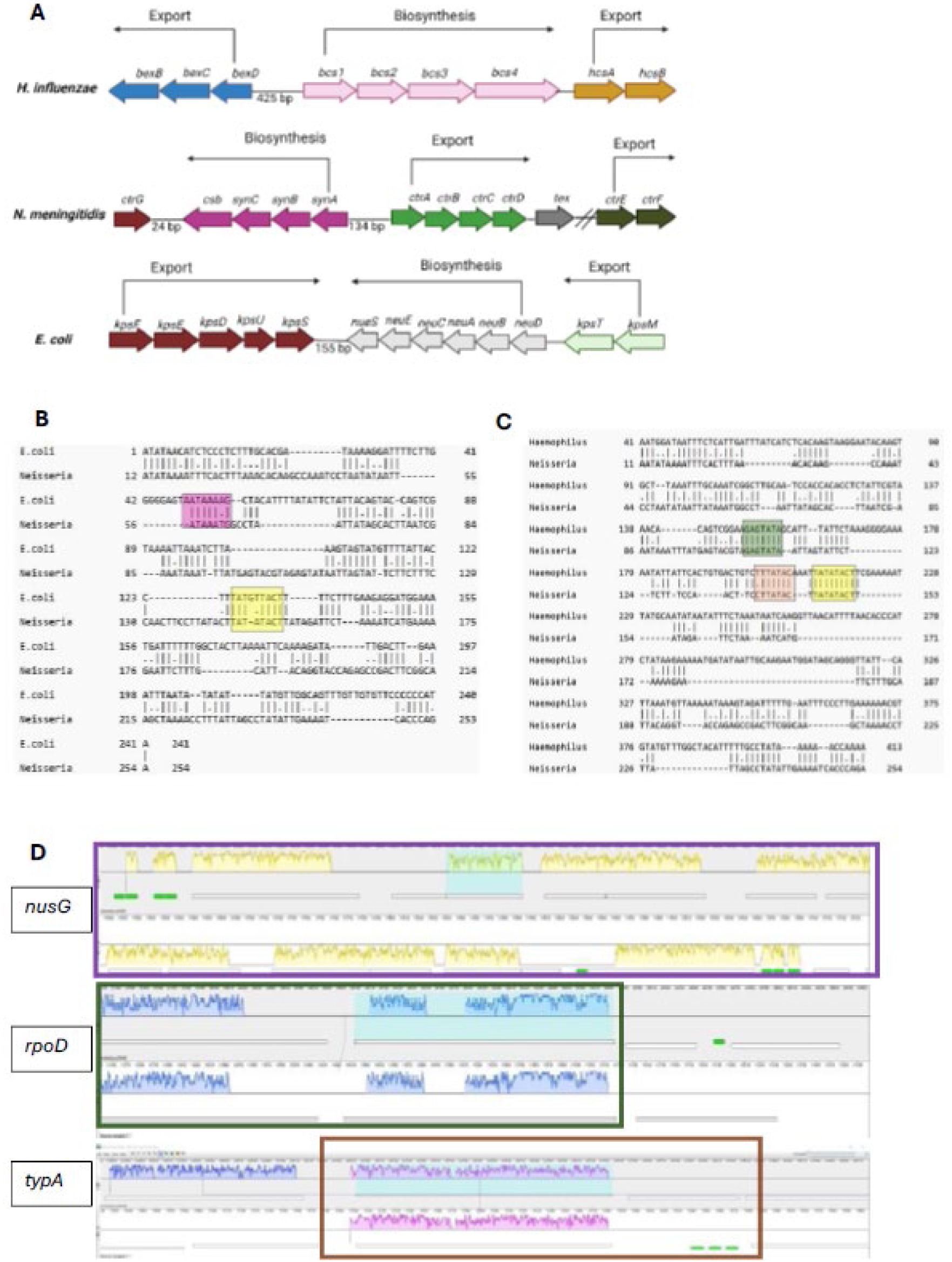
Comparative analysis of Cis-regulatory elements and transcription factors involved in capsule expression in *N. meningitidis* B and related bacteria that use ABC transport-dependent pathways. (A)Genetic organization of the H. influenzae, *E. coli* K1, and *N. meningitidis* capsule loci. The arrows indicate the direction of transcription. An intergenic region separates the start codon of biosynthesis and transport genes. Created in BioRender. Sofowora, I. (2024) BioRender.com/b75z326. (B) Local alignment of the *E. coli* and *N. meningitidis* intergenic regions. Analysis of similar regions revealed conservation in AATAAAAG, highlighted in pink, and TATATACT, highlighted in yellow. These sequences are putative binding sites of phoB and lexA. (c) Local alignment of the H. influenzae and *N. meningitidis* intergenic regions. Analysis of similar regions revealed conservation in GAGTATA, highlighted in green; CTTATAC, highlighted in orange; and TATATACT, highlighted in yellow. These sequences are putative binding sites of rpoD 16, rpoD 18, and lexA. (D) MAUVE alignment of *N. meningitidis* MC58 and *E. coli* K1. The alignment revealed that nusG (highlighted with a purple box), rpoD (highlighted with a green block), and typA (highlighted with an orange box) are in the LCB with several translational proteins and some tRNAs in both Nm and *E. coli*, suggesting their functional importance and evolutionary conservation.

### Identification of conserved transcription factors involved in capsular synthesis in ABC transporter-dependent microbes

Comparative genomic analysis was conducted to elucidate the evolutionary relationships and potential regulatory mechanisms involved in capsule synthesis. BLAST analysis revealed that two of the three TFs (*typA* and *nusG*) required for capsular synthesis in *E. coli* are conserved in *N. meningitidis*, whereas the *mprA* homolog was not detected.

The genome alignment of *N. meningitidis MC58* and *E. coli K1* via the Mauve alignment tool revealed that *the nusG, rpoD,* and *typA* genes are in an LCB with several translational proteins and some tRNAs (colored in green) in both organisms (Fig. 5). This LCB visualized by colored blocks in the alignment highlights areas of high sequence similarity and evolutionary conservation. The co-localization of these TFs within the LCB supports the hypothesis that the capsular genes in both organisms are functionally related and are inherited as a cluster through horizontal gene transfer. The association with translational proteins and tRNAs further suggests that the capsular genes are translationally regulated.

### Trans-complementation assay

The addition of *nusG* and *misR/misS* to the NmIR resulted in a reduction in GFP expression (Table 3). The addition of *typA, crgA, misR,* and *misS* did not affect capsular expression, whereas the addition of the helix–turn–helix (HTH) with the *lexA*-like domain led to an increase in expression (Fig. 6).

**Table 3.** Transcription factors regulating the *synABCD* operon promoter. Transcription factors with the potential to regulate synABCD operon were cloned into a pET22b compatible plasmid, plasmid pstV28, and co-transformed NmIR-GFP-pET22b into E. coli DH5α. Fluorescence intensity was measured with confocal microscopy. Image analysis was conducted with ImageJ

| TFs | Mean Fluorescence<br>Intensity (arbitrary units) | % WT | Standard<br>deviation | P-value |
| --- | --- | --- | --- | --- |
| WT | 3.95 | 100 | 0.78 |  |
| *crgA* | 4.04 | 102.44 | 0.75 | >0.9999 |
| *misR* | 4.06 | 102.99 | 0.72 | >0.9999 |
| *misS* | 3.08 | 78.14 | 0.54 | 0.4838 |
| *misR/misS* | 2.40 | 60.89 | 0.29 | 0.0328 |
| *nusG* | 2.07 | 52.33 | 0.50 | 0.005 |
| *typA* | 4.32 | 109.35 | 0.56 | 0.9859 |
| *HTH_XRE* | 4.88 | 123.75 | 2.25 | 0.5162 |
| *plus ΔlexA* |  |  |  |  |
| *HTH_XRE* | 6.80 | 172.39 | 4.36 | <0.0001 |

**Figure 6.**
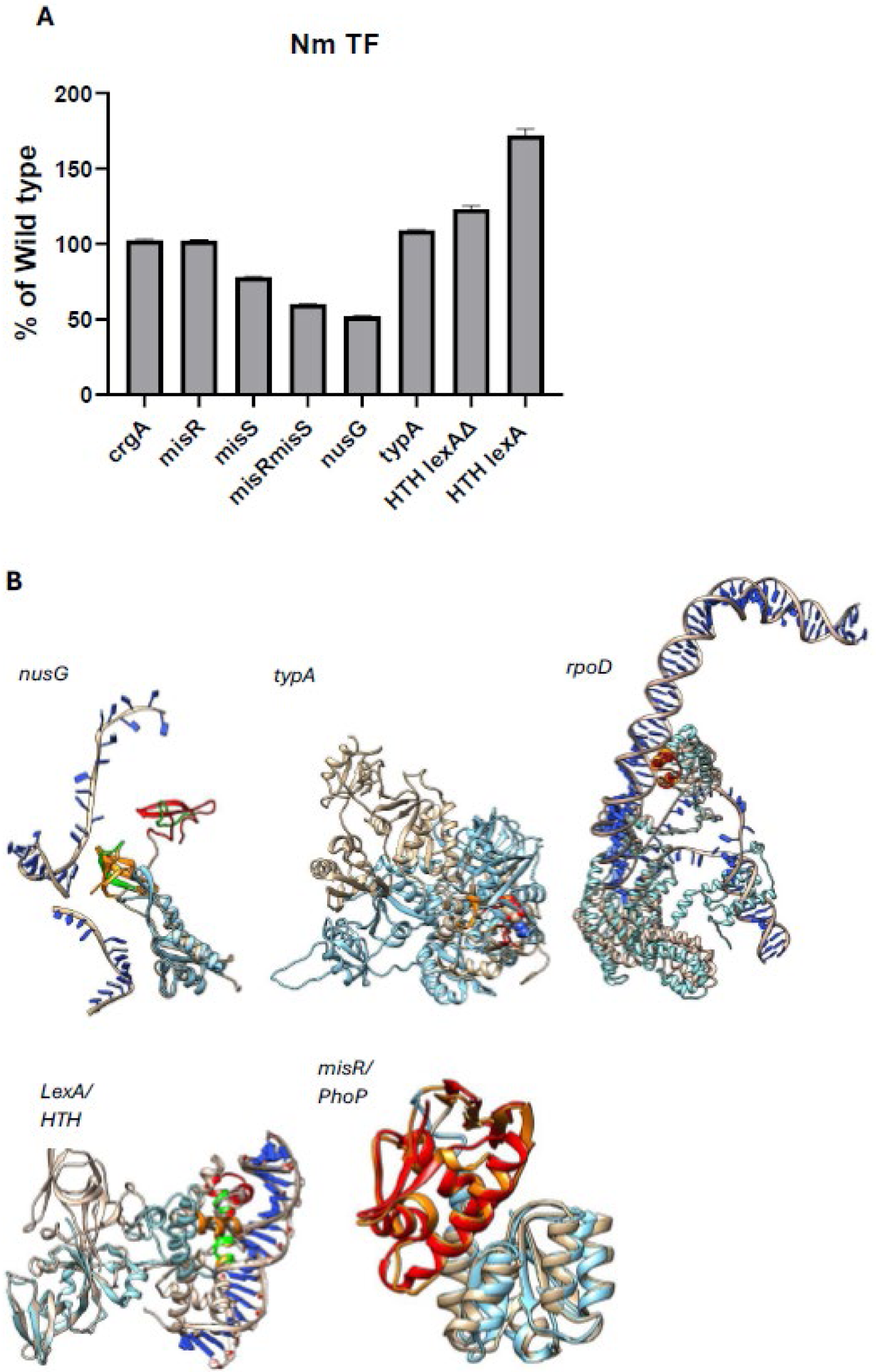
Structural and trans-complementation assay of Transcription factors involved in capsule regulation. (A) Effects of TFs on capsule expression in *N. meningitidis*. The graph shows the relative fluorescence levels (% WT) compared with those of NmIR (WT). The addition of the HTH regulator led to increased capsular expression in the NmIR template. In contrast, the addition of HTH to the template without the lexA binding site did not affect expression. MisR/misS and nusG reduced GFP expression, indicating their role as repressors. In contrast, crgA, misR, misS, misR/misS, and typA had no significant effect on GFP fluorescence, suggesting their roles as activators. (B) Structural alignment of TFs showing conservation of the binding motif. The blue structures represent *N. meningitidis* TFs, whereas the gold structures represent *E. coli* TFs. The binding motif of the *N. meningitidis* TF is colored orange, whereas the binding motif of *E. coli* is colored red. The sequences colored in green represent similarities in the non-conserved regions. The structural alignment shows conservation in the DNA domains of the TFs (lexA/HTH, misR/phoP, rpoD, and typA), suggesting a shared functional role in capsule regulation. In contrast, nusG shows divergence but shares conserved DNA binding motifs, suggesting functional overlap despite structural differences. PDB and AlphaFold IDs: *E. coli* (Eco) nusG: 5MS0 chain F; *N. meningitidis* (Nm) nusG: AF-P65592-F1; Eco typA: 5A9X chain A; Nm typA: AFQ9JZB7-F1; Eco rpoD: 3IYD chain F; Nm rpoD: AF-A0A0H5DMB2-F1; Eco lexA: 3jsp chain A; Nm HTH_XRE: AF-Q9JZT1-F1-v4; Eco phoP: AF-P23836-F1 and Nm misR: NZCP021520 (predicted with Neurosnap AlphaFold2) [18,19].

### Conservation of DNA-Binding Motifs in Transcription Factors Regulating Capsule Production in ABC Transporter-Dependent Organisms**”**

Multiple sequence alignment revealed a high degree of conservation in the DNA binding motifs of the TFS involved in capsule regulation across ABC transporter-dependent organisms. The conservation of the binding motif of *typA* and *nusG* suggests their critical role in ABC transporter-dependent organisms, including *Streptococcus pneumoniae, N. meningitidis, E. coli, and Haemophilus influenzae*. The alignment further highlights the evolutionary relationships between *N. meningitidis and E. coli*, as evidenced by the conserved motifs in *the lexA* and HTH regulators, *rpoD*, and *misR/misS* and its homolog in *E. coli phoP/phoQ.* Structural alignments revealed high structural homology for *lexA, misR, phoP, rpoD,* and *typA,* further supporting the shared functional role of the TFs in these organisms (Fig. 6). In contrast, *nusG* was moderately homologous.

## Discussion

*N. meningitidis* serogroups B, C, Y, and W express sialic acid-containing capsules that are synthesized by transport and synthesis operons that are separated by a 134 bp central region that harbors the respective promoters (5). Of note, the operons are transcribed in opposite directions with the transcription start sites being separated by 1 bp, the promoter for the transport operon is devoid of a -35 motif, and a direct repeat sequence in the 5’-UTR of the synthesis operon is predicted to form a stable hairpin that is postulated to regulate translation by sequestering the ribosome binding site (5). In addition, the region contains regulatory elements and an a MisR binding site but the precise TFBS has not been identified (5). This study investigated the transcriptional regulation of capsular expression in *N. meningitidis B* MC58 (ATCC 53415D-5), with a focus on transcriptional control. Multiple Potential binding sites for the TFs Fis and LexA and three for sigma^70^ (*rpoD*) were predicted using BPROM software that uses linear discriminant function (14). A promoter associated with these TFBSs was predicted with the TSS coinciding with the “A” of the initiation codon implying a leaderless mRNA. Leaderless mRNAs are present in all domains of life and in bacteria they can comprise a significant number of mRNAs, for example, 34% in some *Firmicutes,* 17% in some *Alphaproteobacteria*, and 8% in some *Gammaproteobacteria* (19). Leaderless mRNAs are implicated in adaptation to stress (19).

Surprisingly, the known promoter the was not detected although a 249 bp fragment encompassing sections of the coding regions of *ctraA* and *synA* (*synX*) was used in the prediction. However, BPROM has 80% specificity and sensitivity for *E. coli* sigma70 promoters (14). The known promoter was mutated in other studies and shown to reduce transcription by twofold (7), suggesting the presence of an additional promoter. Thus, systematic deletion using site directed mutagenesis was undertaken in this study.

To aid in the study of capsular expression under different conditions and growth phases, an NmIR-GFP construct was created via directional cloning. NmIR was shown to be active in *E. coli* with increased GFP expression at the stationary phase (Fig. 2), which is consistent with the published literature (7, 17, 20). Indeed, upregulation of capsule expression based on growth phase is seen in other bacteria (21). The deletion of the identified UP-like element preceding the -35-box resulted in a twofold reduction in capsular expression, and further deletion to include the -35 box did not exacerbate this effect. This finding is consistent with the study of von Loewenich et al. (2001), who showed that the deletion of the UP element reduced expression by 50%, but further deletion of the -35 and -10 boxes did not lead to further reduction in reporter-gene expression (7). In contrast, this study revealed that deleting the -10 box alone led to a 57% decrease in expression, highlighting the significant role of the -10 promoter in transcription initiation (Table 1). The NmIR fragment is predicted to contain three *rpoD* sites (Table 1), and the deletion of two of them (*rpoD 16 and rpoD* 17) resulted in a more than 50% reduction in capsular expression, suggesting a function in capsule synthesis. The predicted rpoD17 site coincides with the -10 region of the putative second promoter (Figure 1). The genome of MC58 contains three sigma factors (*rpoD*, *rpoH,* and *rpoN*) (23) and only RpoD binding sites are predicted in the *cps* locus intergenic region.

RpoD (σ^70^) is a conserved sigma factor controls transcription in bacteria during the exponential growth phase (24). A study by Yin et al. (2013) demonstrated that the competition between σ^70^ and other sigma factors for binding sites to RNA polymerase is important for regulating alginate production, with the alternative, stress-related sigma factors leading to enhanced production this capsular polysaccharide in *Pseudomonas aeruginosa* (26). The use of dual promoters in capsule expression has also been reported in *Staphylococcus aureus* whereby one dominant promoter under the control of SigB sigma factor is located downstream of a SigA dependent promoter and the two promoters are regulated by different mechanisms (21). While parallels cannot be drawn at this time, it is known that *S. aureus* capsule production is subject to regulation by complex interactions between the SaeRS bacteria two-component regulatory system (TCS) and the transcription factors that act on competing promoters (27, 28). Similarly, alginate production in *P. aeruginosa* is regulated by the KinB/AlgB TCS with KinB being the sensor kinase and AlgB being the transcription regulator that influences promoter function (29). The TCS is also crucial to capsule expression in *N. meningitidis* capsule expression with MisS functioning as the sensor kinase for the regulator MisR (30). While MisR binds directly to DNA, interaction with specific promoter elements has not been reported. Indeed, sequence analysis identified the potential MisR binding site within the *syn*A coding region (Figure 1B). In these bacteria, the TCS appears to exert negative regulation, which is confirmed in this study for MisS/MisR (Table 3).

Bartley et al. (2013) reported the binding of MisR to NmIR and noted the upregulation of both the *ctrA* and *synA* genes in *misR* deletion mutants (10). In another study, the upregulation of caspule expression was reported in *misR*/*misS* mutants that was accompanied by upregulation of *ctrD*, the gene for the capsule export ATP-binding protein (12). The co-transformation of the *misR/misS* genes with NmIR reduced GFP expression, indicating the negative regulatory function of MisR in capsule expression.

The conservation of the *lexA* binding sites in ABC transporter-dependent *cps* loci despite the absence of the *lexA* gene in *N. meningitidis,* as previously reported by Davidsen & Tønjum (2006), suggests functional significance (Figure 5) (33). Deleting this site led to a 47% increase in capsule expression, suggesting a potential regulatory role of a *lexA–like* factor. Thus, a BLAST search of the MC58 genome was conducted with the *E. coli lexA* gene sequence, and HTH_XRE was identified as a homolog. Cloning of HTH_XRE and subsequent trans-complementation with NmIR-GFP led to a 72% increase in capsule expression. In contrast, no change in expression was observed when the *lexA* site was absent, indicating the specificity of this interaction. This result highlights a novel regulatory mechanism. Further studies are needed to characterize the regulatory network involving the HTH_XRE and its potential impact on adhesion or pathogenesis.

Other TFs studied were Fis, because of the identified binding site (Table 1), CrgA, which has been implicated in the downregulation of capsule during adhesion to epithelial cells, and TypA and NusG because of their implication in capsule expression from analogous *E. coli* operons (8, 34). Fis (a factor for inversion stimulation) is a small basic protein that is important in bacterial nucleoid assembly and recombination processes, and it is also involved in the regulation of many genes, including those involved in two-component systems and lipopolysaccharide biosynthesis (35, 36). The deletion of the repeat region, which spans the *fis* binding site, did not affect capsule expression, as measured by GFP fluorescence. Similarly, *CrgA*, a LysR-type transcriptional regulator, did not influence promoter activity, which is consistent with the findings of von Loewenich et al. (2001). This result notwithstanding, a CrgA binding site that overlaps the -35 promoter region has been identified (8). The caveat to these results is that they were conducted in *E. coli,* and the regulation by *crgA* and *misR* appears to be related to *N. meningitidis* adhesion to epithelial cells (8, 9).

In *E. coli*, three TFs (NusG, TypA, and MprA) are important for the transcriptional control of capsule synthesis (37). Comparative analysis revealed that the loci for the *nusG* and *typA* genes, including adjacent sequences, are conserved in *E. coli* and *N. meningitis* (Fig. 5). Both *N. meningitis nusG* and *typA* were cloned and used in trans-complementation studies. NusG is a universally conserved TF that binds to RNA polymerase to drive RNA synthesis and influence Rho-dependent termination (38). It is known to function as both a terminator and an anti-terminator (38, 39). The observed reduction in capsule expression suggests that in the *N. meningitidis* capsule synthesis operon, NusG primarily acts as a transcription terminator. There was no significant effect of *typA* on GFP expression upon trans-complementation with NmIR. Structural alignment between available structures and Alphafold models show the conservation of the DNA-binding regions for LexA/HTH_XRE, MisR/PhoP, NusG, RpoD, and TypA across species indicating their critical role in protein function and DNA interactions (Figure 6).

## Conclusion

### A GFP fluorescence reporter plasmid was used to further delineate the mechanisms regulating

*N. meningitidis* capsule expression promoter. Novel TFBSs were identified thereby extending the known mechanisms that involve MisR/MisS and CrgA. HTH_XRE, a *N. meningitidis* gene with unknown function was identified as a potential factor regulating capsule production. This factor potentially regulates the downstream promoter for leaderless mRNA. Leaderless mRNAs correlate with stress and results presented in this study indicate the stimulation of GFP expression during stationary phase. Therefore, similar to other bacteria pathogens, the regulation of capsule expression appears to involve multiple mechanisms and TFs. These studies will have utility in enhancing the production of capsular polysaccharides-based biomaterials.

## Materials and methods

### Bioinformatic analysis

The *N. meningitidis* serogroup B sequence was retrieved from the National Center for Biotechnology Information (NCBI) gene bank (X78068.1). The promoter region and putative transcription factor binding sites were predicted via BPROM from Softberry (14). The Basic Local Alignment Search Tool (BLAST) was used to identify homologous genes and conserved motifs.

Jalview was used to visualize the multiple sequence alignments (40). A web logo was generated to represent the conserved residue in a graphical format (41). Mauve software, a multiple genome alignment tool was used to determine the locally conserved blocks (LCBs) in the respective chromosomal regions (42). The binding sites of the TFs in the intergenic region were mapped via FIMO software (43).

### Construction of the reporter strain with *Nm* intergenic region (*NmIR*) activity

The NmIR was PCR amplified by designing primers that incorporated restriction enzyme sites BamH1 (5’-TTT <u>GGA TCC</u> GTG ACG AAT ATA AAA TTT CAC TTT-3’ and NCOI (5’-A CA<u>C CAT GG</u>T ATT TTC AAT ATA GGC TAA TAA AGG TTT TAG CTT -3’) at the ends, allowing the directional cloning of the PCR product into the plasmid expression vector pRSET-EmGFP (Thermo Fisher Scientific, Waltham, MA). The cassette containing NmIR fused to EmGFP was then transferred to the *Bam*H1/*Hin*dIII site of PET 22b, which had been double digested with *Bam*H1 and *Hin*dIII restriction enzymes (Invitrogen). The digested vector and PCR product were ligated and transformed into competent *E. coli* DH5α cells, yielding NmIR-GFP. The desired plasmid constructs were identified via a panel of colony PCRs and confirmed through Sanger sequencing.

### Construction of NmIR mutants and GFP fluorescence measurement

An NEBQ5 mutagenesis kit was used to create deletions of the identified binding sites and the TFs in the NmIR. NEB base changer v1; primer design software was used to design primer pairs for the respective TFs and binding sites. The mutations and primers are listed in Supplementary Table 1, with the deleted sequence in bold. The different regions were amplified, yielding a linear product missing the region deleted. The enzyme *DpnI* (KLD) was used to digest the template, and the linear product was ligated to a circular version, which was subsequently transformed into competent *E. coli* DH5α cells. Plasmid DNA was isolated from selected colonies and sequenced via Sanger sequencing at the Genetic Resources Core Facility at Johns Hopkins University. Capsule expression was measured by quantifying the GFP intensity via confocal microscopy and the ImageJ region of interest manager (ROI) tool (44). Five replicates of confocal images per experiment were imported into image J for intensity analysis. Using ROI manager a minimum of 2000 cells were analyzed.

### TF reporter assay

To quantify and determine the effects of the TFs on capsule synthesis, the sequences for *crgA, lexA, misR, misS, nusG,* and *typA* were retrieved from GenBank with accession number AE002098.2. The respective TF was amplified via primers designed with a ribosome binding site upstream of the ATG initiation codon via the Takara In-Fusion Snap primer design tool. The primers used are listed in Supplementary Table 2. The amplified products were purified (using the NEB Monarch PCR Cleanup Kit) for directional cloning into the BamH1 site of the pstV28 vector via the Takara cloning kit according to the manufacturer’s recommended procedure. The cloned genes were confirmed via Sanger sequencing.

### Trans-complementation assay

The genes were transformed into competent NmIR*-GFP-carrying* cells and cells carrying respective deletion mutants of the putative control elements as predicted with software. After confirmation of transformation, the cells were cultured overnight and imaged with a Leica STELLARIS 5 Confocal Microscope (Leica Microsystems Inc., Deerfield, IL).

### Statistical analysis

Statistical analysis was performed on 3 independent experiments. One-way ANOVA and graphing were accomplished using GraphPad prism 10 software. The mean fluorescence intensity measurement using image J was determined from five replicates per experiment.

## Acknowledgments

The authors acknowledge grant support through NSF DMR Award 2100978 and the use of core facilities supported by the National Institute on Minority Health and Health Disparities through grant numbers 5U54MD013376 and 2U54MD013376-06A1 and National Institute of General Medical Sciences through grant number 5UL1GM118973. The authors thank the Genetic Resources Core Facility, RRID: SCR_018669, Johns Hopkins University, for sequencing services.

